# Enolase-1 is Essential for Neutrophil Recruitment During Acute Inflammation

**DOI:** 10.1101/2024.12.26.630451

**Authors:** Hsueh-Yen Lu, Ping-Hsiang Huang, Ting-Wei Lee, Hui-Wen Chang, Nai-Yu Chen, Yu-Jing Zhuang, Ta-Tung Yuan, Chun-Jen Chen

**Author notes:** These authors contributed equally (Ping-Hsiang Huang, Ting-Wei Lee).

## Abstract

Enolase-1 (ENO1) is a moonlighting protein with multiple functions. When expressed on the cell surface, ENO1 binds plasminogen (PLG) and promotes cell migration by facilitating plasmin (PLM)-mediated extracellular matrix degradation. Here, we observed that inflammatory stimulation significantly upregulated ENO1 expression on the neutrophil surface, both in vitro and in vivo. An anti-ENO1 monoclonal antibody (mAb), 7E5, which blocks the ENO1-PLG interaction, effectively suppressed neutrophil transmigration. In mouse models of acute inflammation, including lipopolysaccharide (LPS)-induced lung injury and necrotic cell challenge, 7E5 treatment markedly reduced neutrophil recruitment and neutrophil extracellular trap (NET) formation. Moreover, 7E5 neutralized the immunostimulatory activity of soluble ENO1, which was significantly elevated in circulation during acute inflammation. These findings highlight ENO1 as a key regulator of inflammation and neutrophil infiltration. Targeting ENO1 with antibodies could be a promising strategy to mitigate tissue damage caused by excessive neutrophilic inflammation.

## Introduction

Neutrophils are the first innate immune cells recruited from the bloodstream to sites of microbial infection or tissue injury during acute inflammation. These cells eliminate pathogens through multiple mechanisms, including phagocytosis, release of granular enzymes and reactive oxygen species (ROS), and formation of neutrophil extracellular traps (NETs) ^1^. Additionally, neutrophils secrete cytokines and chemokines to regulate other leukocytes ^2^. However, these antimicrobial responses are inherently cytotoxic and may cause collateral tissue damage when inflammation becomes dysregulated. This can contribute to autoimmune diseases or chronic inflammation ^3–5^. In the context of acute respiratory distress syndrome (ARDS) during the COVID-19 pandemic, severe acute respiratory syndrome coronavirus 2 (SARS-CoV-2) infection can trigger excessive neutrophil recruitment and activation. This leads to lung injury, cytokine storm, septic shock, and even multiple organ failure ^6,7^.

Upon infection or injury, neutrophils migrate from the bloodstream to sites of insult via a complex multistep cascade. Cytokines and chemokines secreted by tissue-resident innate immune cells stimulate neutrophils to express adhesion molecules, such as integrins and selectins, which interact with counter-receptors on endothelial cells, the basement membrane, and pericytes, enabling neutrophil extravasation ^8^. As neutrophils transmigrate across the endothelium and navigate the extracellular matrix (ECM), neutrophil proteases can facilitate cellular movement by cleaving proteins in the vascular endothelium and ECM ^9–11^. Additionally, the plasminogen (PLG)/tissue-type plasminogen activator (tPA)/urokinase-type plasminogen activator (uPA) system promotes neutrophil adhesion, migration, and thrombolysis ^12–14^. In diseases characterized by excessive neutrophilic inflammation, targeting neutrophil recruitment and activation may mitigate tissue damage and facilitate inflammation resolution ^15,16^.

Enolase-1 (ENO1, or α-enolase) is a glycolytic enzyme that catalyzes the conversion of 2-phosphoglycerate to phosphoenolpyruvate and is ubiquitously expressed in most tissues. Interestingly, ENO1 has been detected on the surface of various cell types, including immune cells ^17–20^, cancer cells ^21,22^, and neuronal cells ^23^. On the cell surface, ENO1 functions as a plasminogen receptor (PLGR), facilitating plasmin (PLM) activation and cell migration. This mechanism contributes to cancer metastasis ^22,24^ and immune cell infiltration ^19^. Cell surface ENO1 may also bind other immunostimulatory molecules, such as apolipoprotein B, which promotes monocyte activation ^25^. Additionally, ENO1 serves as an autoantigen, triggering autoimmune responses in diseases such as rheumatoid arthritis (RA) ^26^, systemic sclerosis (SSc) ^27^, systemic lupus erythematosus (SLE) ^28^, and inflammatory bowel disease ^29^. Beyond its role as a PLGR, ENO1 is released from injured cells and acts as a damage-associated molecular pattern (DAMP), activating monocytes, neutrophils, and endothelial cells ^30,31^. ENO1 levels increase in the circulation following acute injury, suggesting a role in regulating inflammatory responses ^31–33^. Thus, ENO1 is a moonlighting protein with diverse immunoregulatory functions that vary based on cellular localization and pathophysiological context.

The PLGR function of ENO1 has been documented on human monocytes ^17^, neutrophils ^18^, and B lymphocytes ^20^. On monocytes, ENO1 surface expression increases upon lipopolysaccharide (LPS) stimulation, promoting monocyte recruitment in an ENO1-dependent manner ^19^. However, despite its known role in PLG activation, the in vivo impact of ENO1 on neutrophilic inflammation remains unclear. Here, we demonstrate that inflammatory stimulation induces ENO1 expression on the neutrophil surface. Furthermore, an anti-ENO1 monoclonal antibody (mAb) effectively attenuates neutrophil recruitment and migration while also neutralizing the DAMP activity of ENO1. Our findings highlight ENO1 as a critical regulator of neutrophil-driven inflammation, suggesting that targeting ENO1 may be a potential therapeutic strategy for controlling excessive neutrophilic inflammation.

## Results

### Surface ENO1 expression on neutrophils is upregulated after LPS stimulation in vitro and in vivo

To investigate surface ENO1 expression on neutrophils during inflammation, we first isolated neutrophils from mouse bone marrow and analyzed ENO1 expression using flow cytometry. Murine bone marrow neutrophils exhibited basal surface ENO1 expression, which significantly increased following LPS stimulation in vitro (Figure 1A). To determine whether surface ENO1 is similarly upregulated in vivo during inflammation, we examined neutrophils from mice challenged intraperitoneally (i.p.) with LPS. Under systemic inflammatory conditions, neutrophils in both the bone marrow and circulation showed elevated surface ENO1 expression (Figure 1B). Likewise, in mice subjected to intratracheal (i.t.) LPS administration to induce acute lung injury (ALI), we observed increased surface ENO1 levels on neutrophils in the bone marrow and blood (Figure 1C). To explore the underlying mechanism of ENO1 upregulation on inflammatory neutrophils, we assessed ENO1 expression at both mRNA and protein levels. Neutrophils from LPS-challenged and control animals expressed comparable levels of ENO1 mRNA (Figure 1D) and total protein (Figure 1E), suggesting that the increased surface ENO1 expression during acute inflammation may result from the exteriorization of cytosolic ENO1 rather than de novo synthesis.

**Figure 1.**
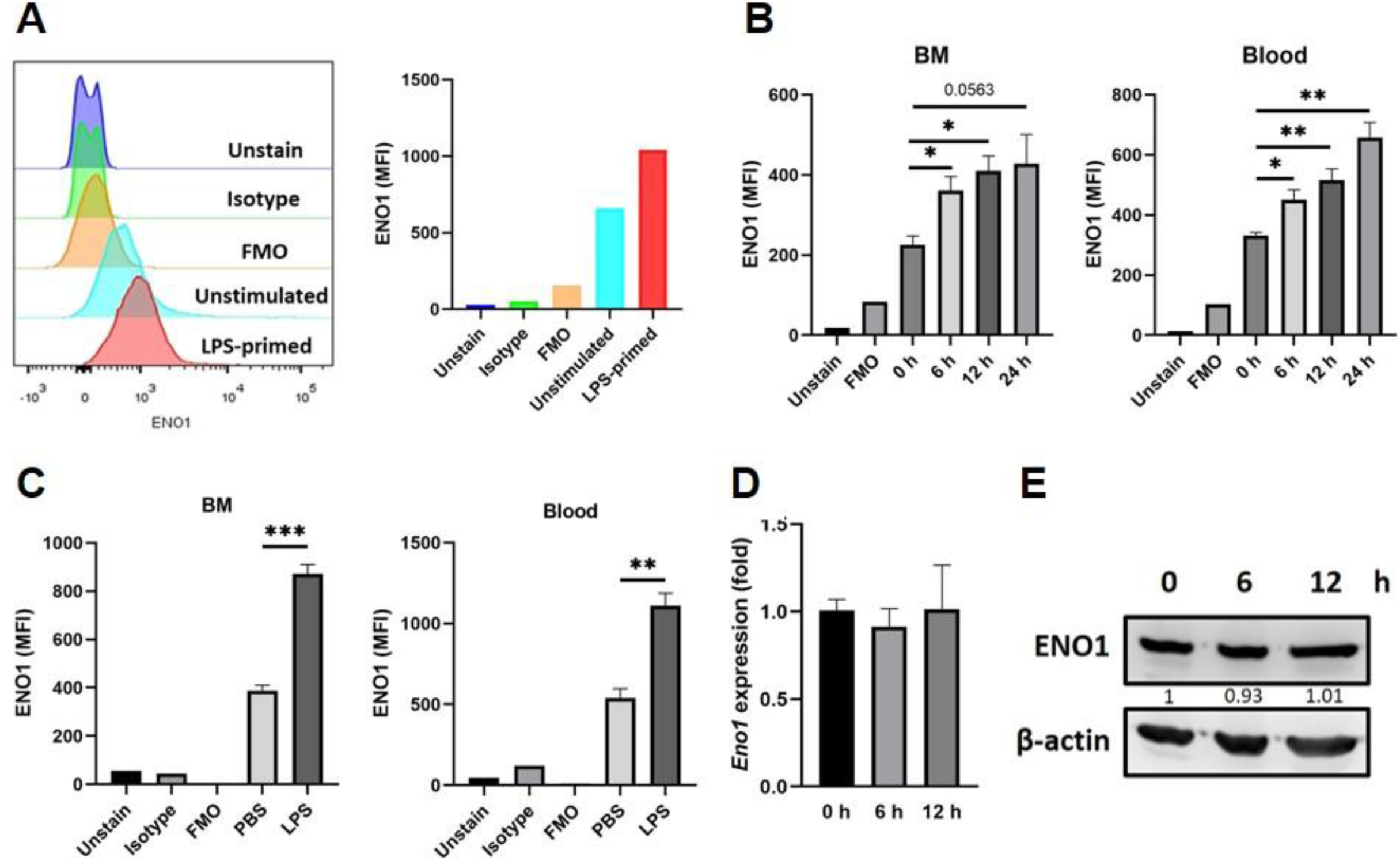
Surface ENO1 expression on neutrophils is upregulated after LPS stimulation in vitro and in vivo. (A) Flow cytometry analysis of surface ENO1 expression on mouse bone marrow neutrophils treated ex vivo with LPS for 3 hours. (B) Flow cytometry analysis of surface ENO1 expression on bone marrow (BM) and blood neutrophils from mice challenged i.p. with LPS for the indicated times (*n* = 3). (C) Flow cytometry analysis of surface ENO1 expression on BM and blood neutrophils from mice 24 hours after i.t. challenge with PBS (*n* = 2) or LPS (*n* = 4). (D) RT-qPCR analysis of ENO1 mRNA expression in neutrophils from mice challenged i.p. with LPS for the indicated times. (E) Western blot analysis of ENO1 protein expression in neutrophils from mice challenged i.p. with LPS for the indicated times. ENO1 signal intensity (normalized to actin) is expressed as fold changes relative to the 0-h group and shown below each lane. Data are representative of two (A, B) or three (C, E) independent experiments. Data in (D) are combined from four independent experiments. Data are presented as mean ± SEM. \**p* < 0.05, \*\**p* < 0.01, \*\*\**p* < 0.001.

### The anti-ENO1 mAb 7E5 disrupts ENO1-PLG interaction and inhibits neutrophil transmigration

ENO1 has been identified as a PLGR that promotes PLM activation on the cell surface ^17–19^. To confirm the interaction between ENO1 and PLG, we conducted a cell-free dot blot assay. As shown in Figure 2A, ENO1 specifically interacted with PLG, while no binding was observed with the control protein, bovine serum albumin (BSA). Next, we evaluated whether the rat anti-mouse ENO1 mAb 7E5 could block the ENO1-PLG interaction using an ELISA-based protein binding assay. Notably, 7E5 mAb significantly disrupted ENO1-PLG binding compared to the control IgG (Figure 2B). To investigate whether ENO1 expression on the neutrophil surface facilitates PLM-mediated neutrophil migration in vitro, we performed a neutrophil transmigration assay. Neutrophils treated with PLG and uPA migrated through an ECM gel-coated membrane in response to the chemokine KC. Notably, pretreatment with mAb 7E5, but not control IgG, inhibited neutrophil transmigration (Figure 2C). As a positive control, neutrophil transmigration was also blocked by tranexamic acid (TXA), a lysine analog that binds PLG and prevents its activation. These findings indicate that ENO1 expression on the neutrophil surface facilitates PLM-mediated proteolysis and neutrophil migration and that this process can be effectively inhibited by the mAb 7E5.

**Figure 2.**
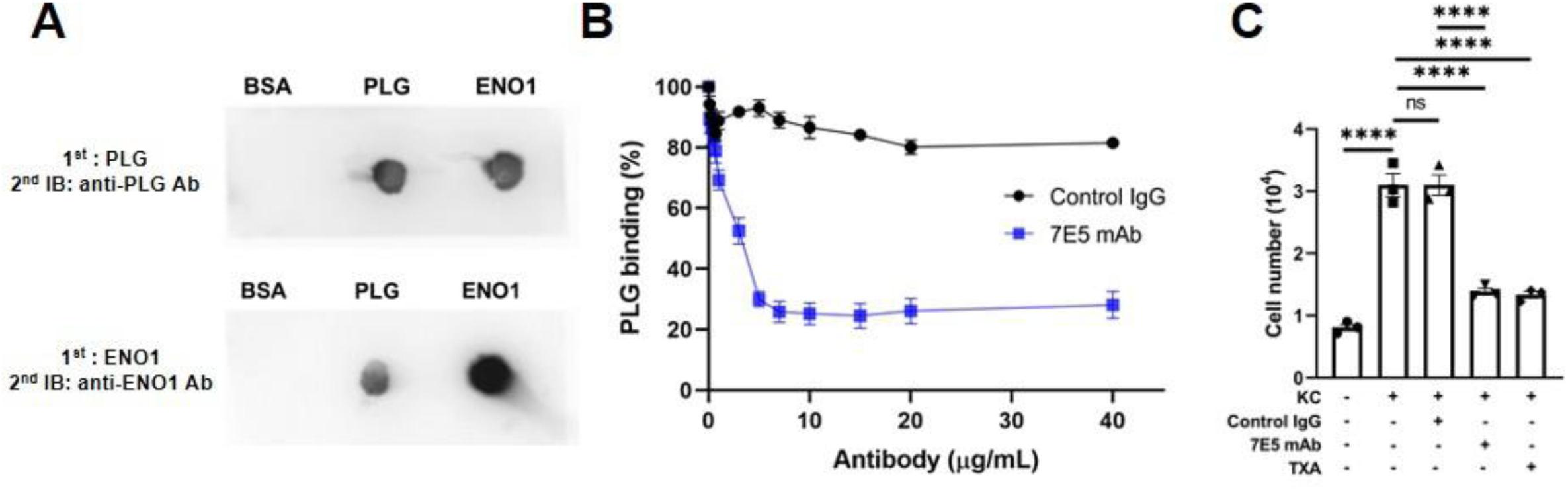
The anti-ENO1 mAb 7E5 disrupts ENO1-PLG interaction and inhibits neutrophil transmigration. (A) Dot blot analysis showing the specific interaction between ENO1 and PLG. (B) ELISA-based protein binding assay measuring PLG binding to plate-bound ENO1 in the presence of 7E5 mAb or control IgG. (C) Neutrophil transmigration assay using cells from naïve mice, assessed in the presence of 7E5 mAb, control IgG, or TXA (*n* = 3). Data in (A) and (C) are representative of two and three independent experiments, respectively. Data in (B) are combined from three independent experiments performed in duplicates. Data are presented as mean ± SEM. \**p* < 0.05, \*\**p* < 0.01, \*\*\**p* < 0.001.

### ENO1 drives neutrophilic inflammation and tissue injury in the LPS-induced ALI mouse model

Given that increased ENO1 expression enhances neutrophil pericellular proteolytic activity, we next investigated its role in neutrophil migration using the LPS-induced ALI mouse model, which is characterized by excessive neutrophil infiltration into the alveolar compartment ^34^. Mice were intravenously (i.v.) administered the anti-ENO1 mAb 7E5 (5 mg/kg) or a control IgG 2 hours before i.t. LPS instillation. The antibody dose was selected based on previous observations that the same dose of 7E5 mAb ameliorated disease severity in a mouse model of experimental autoimmune encephalomyelitis ^35^. At 24 hours post-challenge, mice were sacrificed, and bronchoalveolar lavage fluid (BALF) was analyzed for inflammatory cell infiltration and proinflammatory mediator production. Lung tissue sections were also examined for histopathological changes. LPS-challenged mice exhibited a robust influx of neutrophils into the alveolar space (Figure 3A). Treatment with 7E5 mAb significantly reduced neutrophil infiltration (Figure 3B), proinflammatory cytokine levels (TNF, IL-6, and KC), and protein accumulation in BALF, a marker of lung edema. Histopathological analysis further demonstrated that 7E5 mAb treatment reduced neutrophil accumulation in both alveolar and interstitial spaces and alleviated lung injury (Figure 3D).

**Figure 3.**
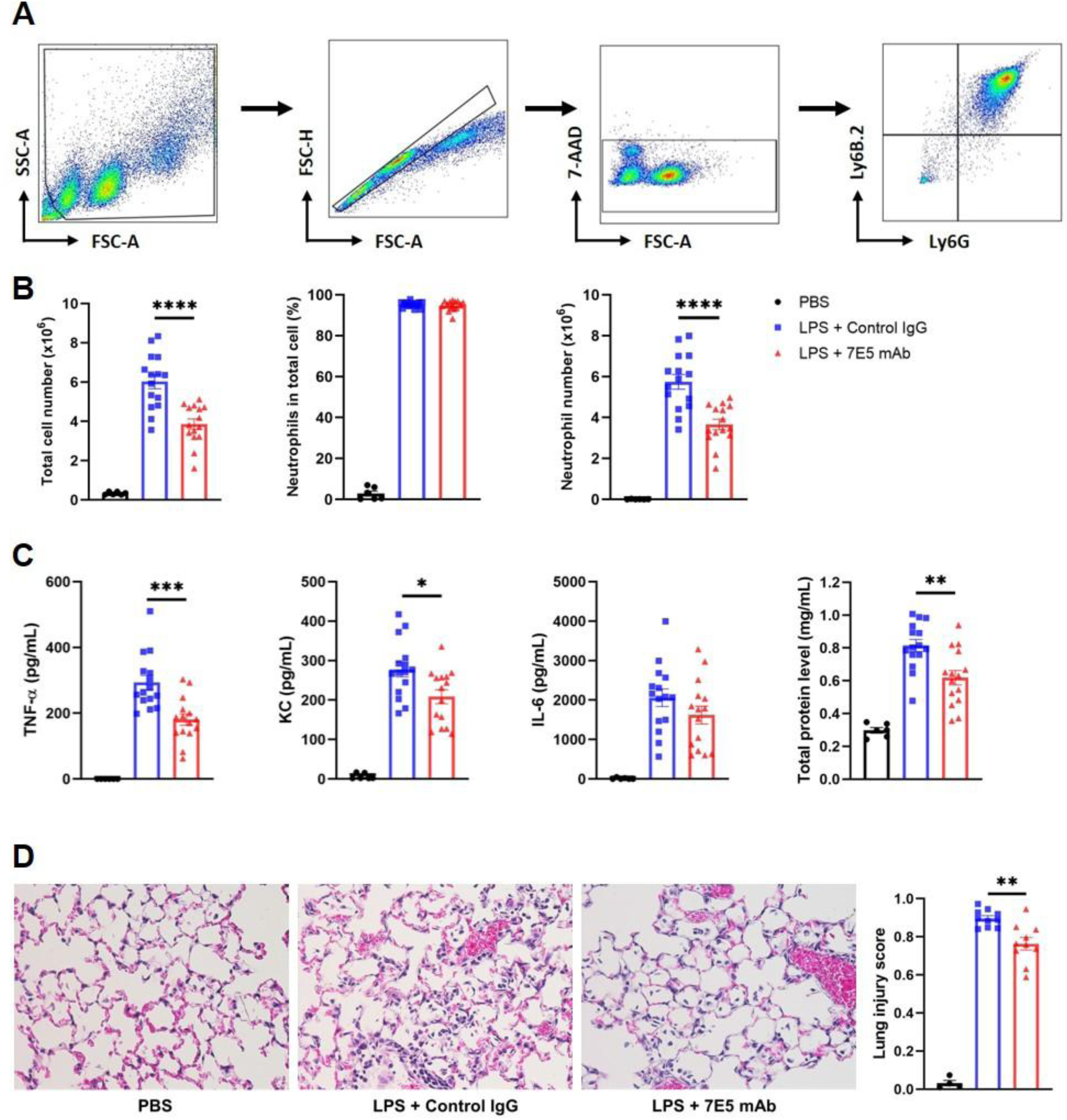
ENO1 drives neutrophilic inflammation and tissue injury in the LPS-induced ALI mouse model. (A) Representative flow cytometry density plots and gating strategy for BALF cells from mice 24 hours after i.t. LPS challenge. Neutrophils are identified as Ly6G^+^Ly6B.2^+^ cells. (B) Total cell counts, neutrophil percentages, and neutrophil counts in BALF from LPS-challenged mice treated with mAb 7E5 or control IgG. Mice receiving i.t. PBS served as the negative control. (C) Cytokine and protein levels in BALF. (D) Representative H&E-stained lung sections and quantitative lung injury scores. Data in (B) and (C) represent combined results from three independent experiments. Data in (D) are from two independent experiments. Data are presented as mean ± SEM. \**p* < 0.05, \*\**p* < 0.01, \*\*\**p* < 0.001, \*\*\*\**p* < 0.0001.

We also investigated the effect of ENO1 blockade on neutrophil extracellular trap (NET) formation, a known contributor to ALI ^36^. Treatment with the 7E5 mAb reduced the number of NETotic neutrophils containing citrullinated histone 3 (citH3) (Figures 4A and 4B) and decreased cell-free double-stranded DNA (dsDNA) levels in BALF (Figure 4C). Additionally, ENO1 blockade lowered the frequency of NETotic neutrophils in the blood (Figure 4B). These findings suggest that ENO1 plays a key role in neutrophilic inflammation and lung injury during LPS-induced ALI.

**Figure 4.**
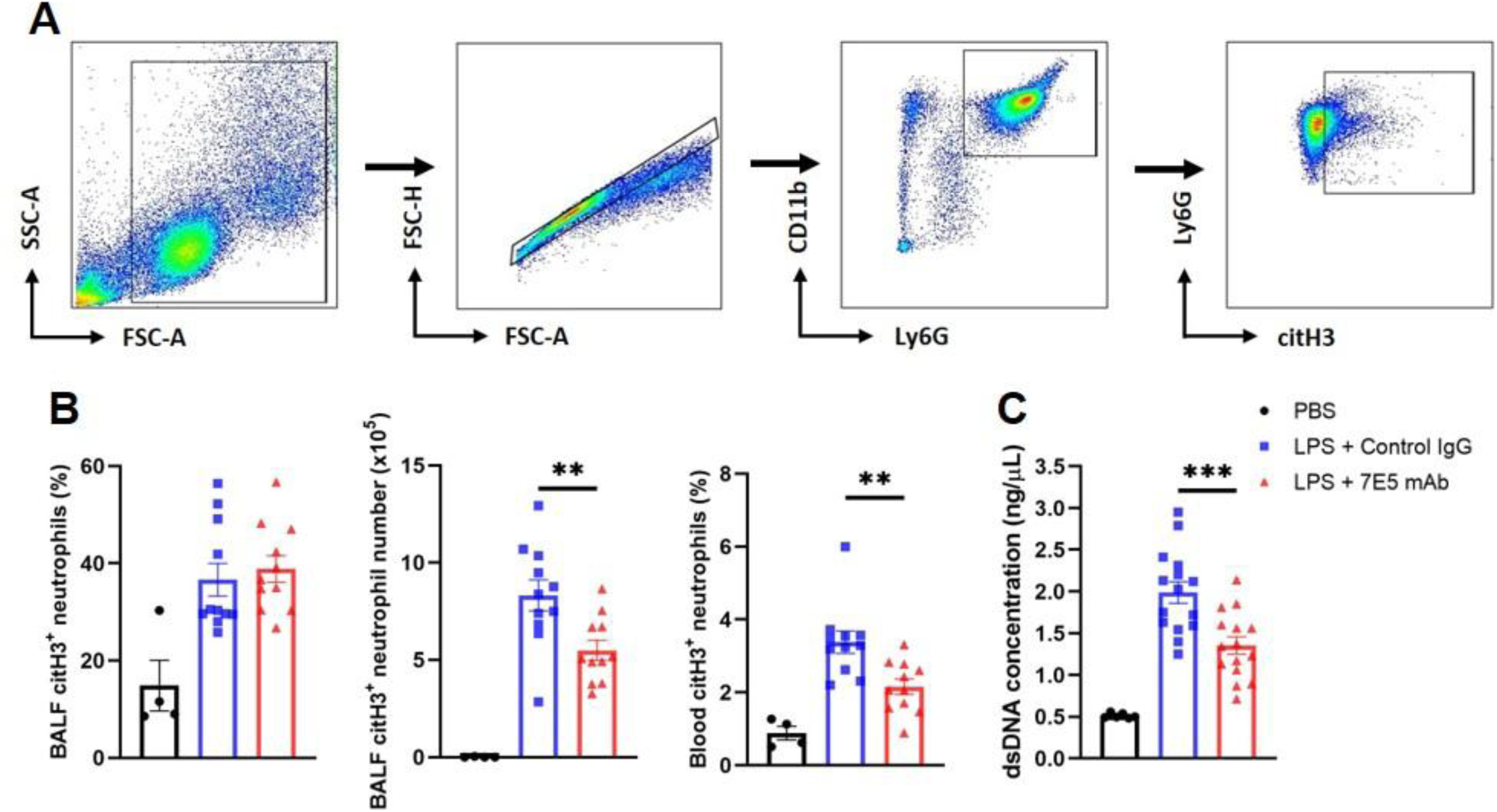
ENO1 blockade reduces NETotic neutrophils in LPS-induced ALI. (A) Representative flow cytometry density plots and gating strategy for Ly6G^+^citH3^+^ NETotic neutrophils in BALF and blood 24 hours after i.t. challenge with PBS, LPS + control IgG, or LPS + 7E5 mAb. (B) Frequencies and absolute counts of NETotic neutrophils in BALF and blood. (C) Cell-free dsDNA levels in BALF. Data in (B) and (C) are combined from two and three independent experiments, respectively, and presented as mean ± SEM. \*\**p* < 0.01, \*\*\**p* < 0.001.

### Anti-ENO1 mAb inhibits neutrophil recruitment in cell injury-induced inflammation

In the absence of infection, acute inflammation can be triggered by DAMPs released from injured cells, with recruited neutrophils potentially exacerbating tissue damage ^37^. To investigate whether ENO1 regulates neutrophil migration during sterile inflammation, we injected necrotic EL4 cells i.p. into mice pretreated with either the 7E5 mAb or control IgG. At 12 hours post-injection, a robust influx of neutrophils and monocytes was observed in the peritoneal cavity (Figure 5A). However, treatment with 7E5 mAb significantly reduced the total number of peritoneal exudate cells (PECs) and diminished neutrophil and monocyte recruitment (Figure 5B). These findings suggest that ENO1 plays a crucial role in neutrophil recruitment to injured tissues during sterile inflammation.

**Figure 5.**
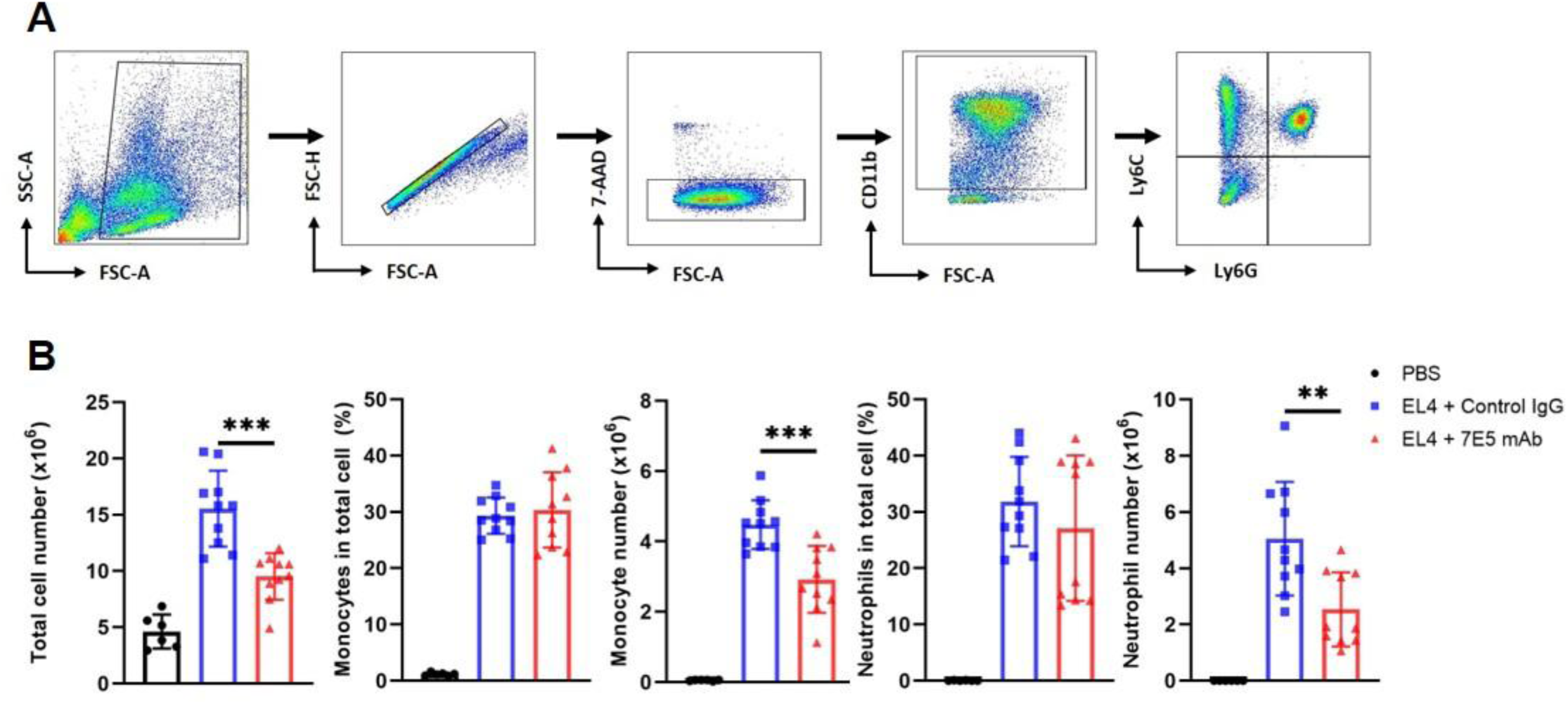
Anti-ENO1 mAb inhibits neutrophil recruitment in cell injury-induced inflammation. (A) Representative flow cytometry density plots and gating strategy for PECs from mice 12 hours after i.p. challenge with necrotic EL4 cells. The CD11b^+^Ly6C^+^Ly6G^-^ and CD11b^+^Ly6C^+^Ly6G^+^ gates represent monocytes and neutrophils, respectively. (B) Frequencies and absolute counts of total PECs, monocytes, and neutrophils recruited to the peritoneal cavity. Data are combined from two independent experiments and presented as mean ± SEM. \*\**p* < 0.01, \*\*\**p* < 0.001.

### 7E5 mAb reduces soluble ENO1-induced alveolar macrophage activation

Elevated levels of ENO1 have been detected in the plasma of post-trauma patients, and soluble ENO1 may directly activate innate immune cells ^30,31^. In both LPS-induced ALI and necrotic cell-induced models of acute inflammation, serum ENO1 levels significantly increased following inflammatory challenges (Figure 6A and 6B). We hypothesized that soluble ENO1 acts as a DAMP, stimulating tissue-resident macrophage activation, and that this effect could be blocked by the 7E5 mAb. To test this, we stimulated alveolar macrophage MH-S cells with recombinant ENO1 protein in the presence or absence of 7E5 mAb. ENO1 stimulation led to a significant increase in TNF and IL-6 production by MH-S cells, while 7E5 mAb dose-dependently suppressed this inflammatory response (Figure 6C). These findings demonstrate that, beyond blocking PLM-mediated neutrophil migration, the anti-ENO1 mAb 7E5 also attenuates inflammation by neutralizing the DAMP activity of soluble ENO1.

**Figure 6.**
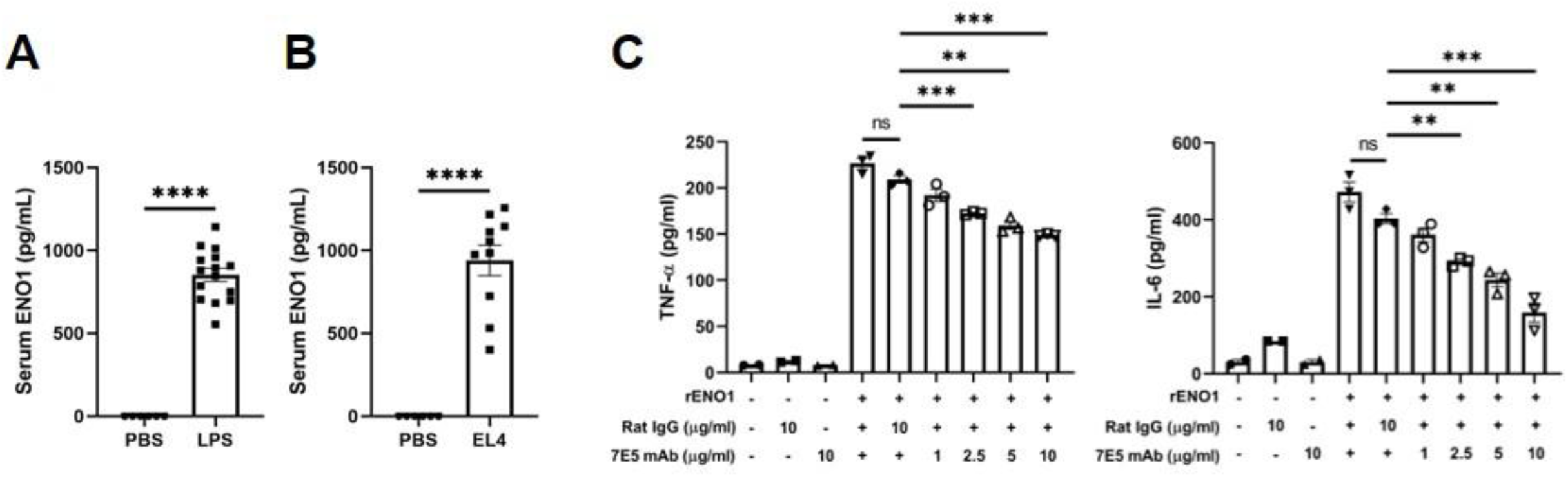
7E5 mAb reduces soluble ENO1-induced alveolar macrophage activation. (A) ELISA analysis of soluble ENO1 levels in plasma from mice 24 hours after i.t. challenge with PBS or LPS. (B) ELISA analysis of soluble ENO1 levels in plasma from mice 12 hours after i.p. challenge with PBS of necrotic EL4 cells. (C) ELISA analysis of TNF and IL-6 production by MH-S cells 20 hours after recombinant rENO1 stimulation, in the presence or absence of 7E5 mAb or control IgG at the specified dose. Data in (A) and (B) are combined from two and three independent experiments, respectively. Data in (C) are representative of three independent experiments. Data are presented as mean ± SEM. \*\**p* < 0.01, \*\*\**p* < 0.001, \*\*\*\**p* < 0.0001.

**Figure 7.**
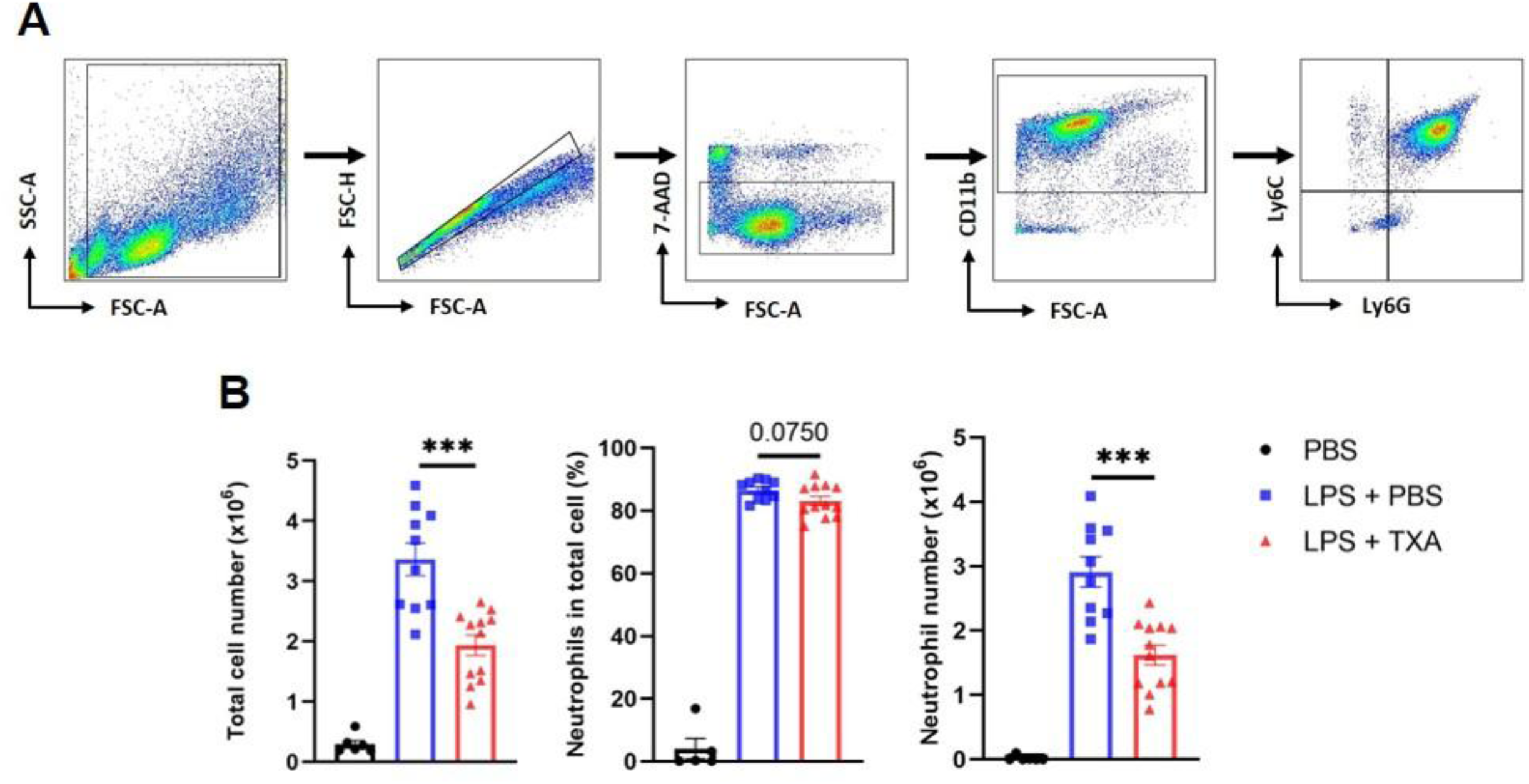
A PLG inhibitor attenuates neutrophil recruitment in LPS-induced ALI. (A) Representative flow cytometry density plots and gating strategy of BALF cells from mice 24 hours after i.t. challenge with LPS. The CD11b^+^ Ly6C^+^Ly6G^+^ gate represents neutrophils. (B) Total cell counts, neutrophil percentages, and neutrophil counts in BALF of LPS-challenged mice treated with PBS or TXA. Mice challenged i.t. with PBS served as negative controls. Data are combined from two independent experiments and presented as mean ± SEM. \*\*\**p* < 0.001.

### A PLG inhibitor attenuates neutrophil recruitment in LPS-induced ALI

To confirm the role of PLG activation in neutrophil migration during inflammation, we examined the effect of TXA, a clinically used PLG inhibitor, on neutrophil recruitment. Mice were i.v. administered PBS or TXA 0.5 hours before i.t. LPS instillation. At 24 hours post-LPS challenge, BALF analysis revealed a robust neutrophil influx in LPS-hallenged mice, which was significantly inhibited by TXA treatment (Figures 6A and 6B). These findings demonstrate that the PLG/PLM system plays a critical role in mediating neutrophil migration during inflammation. Collectively, these findings demonstrate that the PLG/PLM system is essential for neutrophil migration during inflammation.

## Discussion

Acute inflammation is a crucial component of the host immune response; however, excessive neutrophil activation can lead to collateral tissue damage and, in severe cases, organ failure. Identifying key molecular mediators of neutrophilic inflammation is essential for developing targeted therapeutic strategies. In this study, we demonstrate that surface ENO1 expression on neutrophils is upregulated during inflammation and facilitates PLG activation, which enhances neutrophil migration through extracellular matrix proteolysis. Using LPS-induced ALI and necrotic cell-induced peritonitis models, we show that administration of an anti-ENO1 mAb, 7E5, significantly attenuates neutrophilic inflammation. Furthermore, we provide evidence that soluble ENO1, which increases in circulation following inflammatory challenges, acts as a DAMP that amplifies the inflammatory response. These findings suggest that ENO1 plays a key role in mediating neutrophilic inflammation and may be a promising therapeutic target for conditions such as ALI/ARDS.

ENO1 is known to be present on the surface of various immune cells, including neutrophils ^17,18,20^. However, it remains unclear whether ENO1 expression on the neutrophil surface is altered during inflammation. Our findings reveal that inflammatory stimulation, both in vitro and in vivo, leads to increased ENO1 translocation to the neutrophil surface without corresponding changes in total ENO1 mRNA or protein levels. This suggests that ENO1 is mobilized from intracellular stores, a process previously observed in monocytes ^19^ and cancer cells ^38,39^. In human breast cancer MDA-MB-231 cells, Ca^2+^ influx mediated by stromal interaction molecule (STIM) 1 and the calcium release-activated calcium (ORAI) 1 molecule regulates ENO1 exteriorization ^38^. In human lung adenocarcinoma A549 cells, protein arginine methyltransferase 5 (PRMT5) was identified as responsible for ENO1 methylation and cell surface trafficking ^39^. The molecular mechanism underlying ENO1 exteriorization in inflammatory neutrophils has not yet been identified, and whether similar mechanisms observed in cancer cells contribute to ENO1 translocation in neutrophils remains to be investigated.

The PLG/PLM system is known to promote cell migration by degrading extracellular matrix proteins ^19,22,40^. Our study demonstrates that neutrophil transmigration is blocked by the 7E5 mAb, which specifically interrupts the interaction between ENO1 and PLG. In the LPS-induced ALI model, neutrophil recruitment was significantly reduced by both 7E5 mAb and the PLG inhibitor TXA, underscoring the essential role of PLM-mediated proteolysis in neutrophil migration. Previous studies have presented differing perspectives on the role of PLG/PLM in neutrophil migration using PLM-deficient mice. Ploplis et al. reported that Plg (+/+) and Plg (-/-) mice showed similar neutrophil recruitment during thioglycolate-induced peritoneal inflammation ^41^. However, in a biomaterial-induced peritonitis model, neutrophil recruitment was markedly reduced in Plg (-/-) mice ^42^. Neutrophil infiltration in response to ischemia/reperfusion injury was also diminished in tPA (-/-) mice ^12^. These discrepant observations may result from the use of different inflammation models in those studies. In the LPS-induced ALI and necrotic cell-induced peritonitis mouse models, our pharmacological inhibition approach strongly supports the contribution of PLG activation to neutrophil transmigration during inflammation.

We also observed a marked increase in circulating ENO1 following inflammatory challenges, suggesting its role as a soluble proinflammatory mediator. In vitro, recombinant ENO1 robustly activated alveolar macrophages, promoting the release of TNF and IL-6, an effect that was dose-dependently inhibited by 7E5 mAb. These findings indicate that ENO1 functions as a DAMP, stimulating innate immune activation and amplifying the inflammatory response. Similar proinflammatory effects of ENO1 have been reported in autoimmune diseases and trauma-induced ALI, where ENO1 activates monocytes via the CD14-TLR4 pathway and stimulates pulmonary endothelial cells and neutrophils through PLM activation of protease-activated receptor 2 ^27,28,30,31^. Given these dual functions of ENO1—both as a PLGR facilitating neutrophil migration and as a soluble immune activator—therapeutic targeting of ENO1 may provide a more comprehensive anti-inflammatory strategy than inhibiting PLG activation alone.

In conclusion, our findings highlight ENO1 as a key mediator of neutrophilic inflammation, promoting both neutrophil migration and immune activation. The anti-ENO1 mAb 7E5 effectively attenuates neutrophil recruitment, limits tissue damage, and neutralizes the proinflammatory effects of soluble ENO1. Given that 7E5 is a rat anti-mouse ENO1 mAb designed as a surrogate for HL217, a humanized anti-human ENO1 mAb ^43,44^, our results provide a strong rationale for further evaluating HL217 as a potential therapeutic for inflammatory diseases characterized by excessive neutrophilic responses.

### Limitations of the study

Our data highlight the critical role of ENO1 in mediating neutrophilic inflammation using mouse neutrophils and acute inflammation models. However, whether this mechanism similarly regulates neutrophilic inflammation in humans remains to be determined. The process by which ENO1 translocates from the cytosol to the neutrophil surface also requires further investigation. In this study, we primarily used antibody-mediated blockade and neutralization to examine the role of ENO1 in inflammation. Due to the lack of suitable neutrophil cell lines, we were unable to use genetic approaches to study ENO1 in neutrophil migration, as previously done in monocytes ^19^. Additionally, since ENO1 is essential for glycolysis, its role in inflammation could not be assessed in animals lacking ENO1 expression.

## RESOURCE AVAILABILITY

### Lead contact

Further information and requests for resources and reagents should be directed to and will be fulfilled by the lead contact, Chun-Jen Chen

### Materials availability

This study did not generate new unique reagents.

### Data and code availability

● This paper does not report the original code.
● This paper does not report nucleotide sequencing-associated datasets, proteomics, peptidomics, metabolomics, structures of biological macromolecules, or small-molecule crystallography.
● Any additional information about the data reported in this paper is available from the lead contact upon request.

## ACKNOWLEDGMENTS

This research was funded by National Taiwan University (109L104703-4, 111L893404, 112L892204, 113L890404). We thank HuniLife Biotechnology Inc. for providing proprietary anti-ENO1 mAbs 7E5 and HL217, and plasmid for recombinant ENO1 protein expression. We thank NTU TechComm flow cytometry and qPCR core facilities.

## AUTHOR CONTRIBUTIONS

Conceptualization: H.-Y.L., P.-H.H., N.-Y.C., Y.-J.Z. and C.-J.C.; methodology: H.-Y.L., P.-H.H., N.-Y.C., Y.-J.Z. and C.-J.C.; validation: H.-Y.L., P.-H.H., H.-W.C. and C.-J.C.; formal analysis: H.-Y.L., P.-H.H., and C.-J.C.; investigation: H.-Y.L., P.-H. H., T.-W.L., and H.-W.C.; resources: H.-Y.L., P.-H.H., T.-T.Y., and C.-J.C.; data curation: H.-Y.L and C.-J.C.; writing – original draft: H.-Y.L and C.-J.C.; writing – review and editing: H.-Y.L and C.-J.C.; visualization: H.-Y.L and C.-J.C.; supervision: C.-J.C.; project administration: C.-J.C.; funding acquisition: C.-J.C.

## DECLARATION OF GENERATIVE AI AND AI-ASSISTED TECHNOLOGIES IN THE WRITING PROCESS

During the preparation of this work, the authors used ChatGPT to enhance language and readability. They subsequently reviewed and edited the content as necessary and take full responsibility for the final publication.

## STAR METHODS

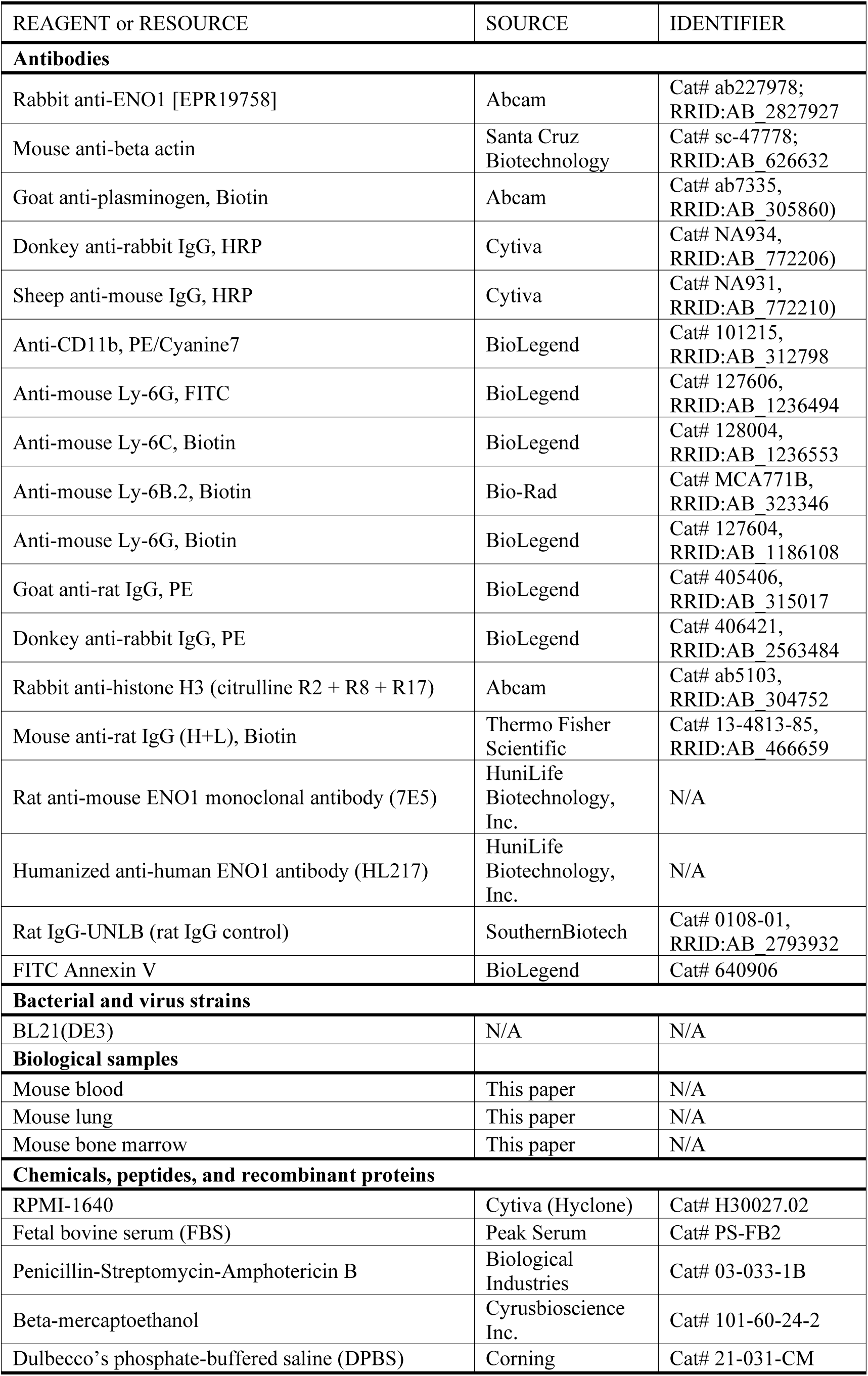
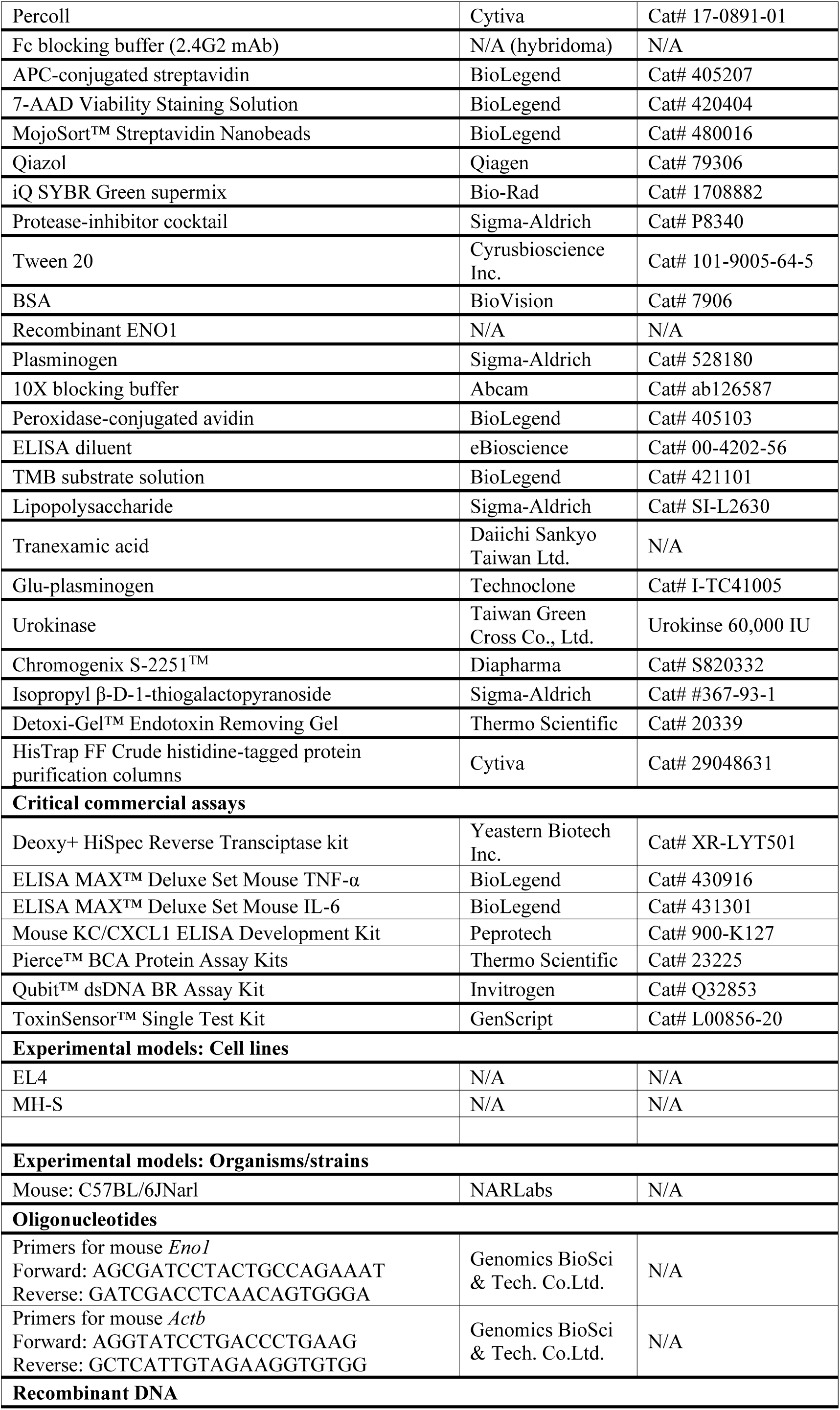
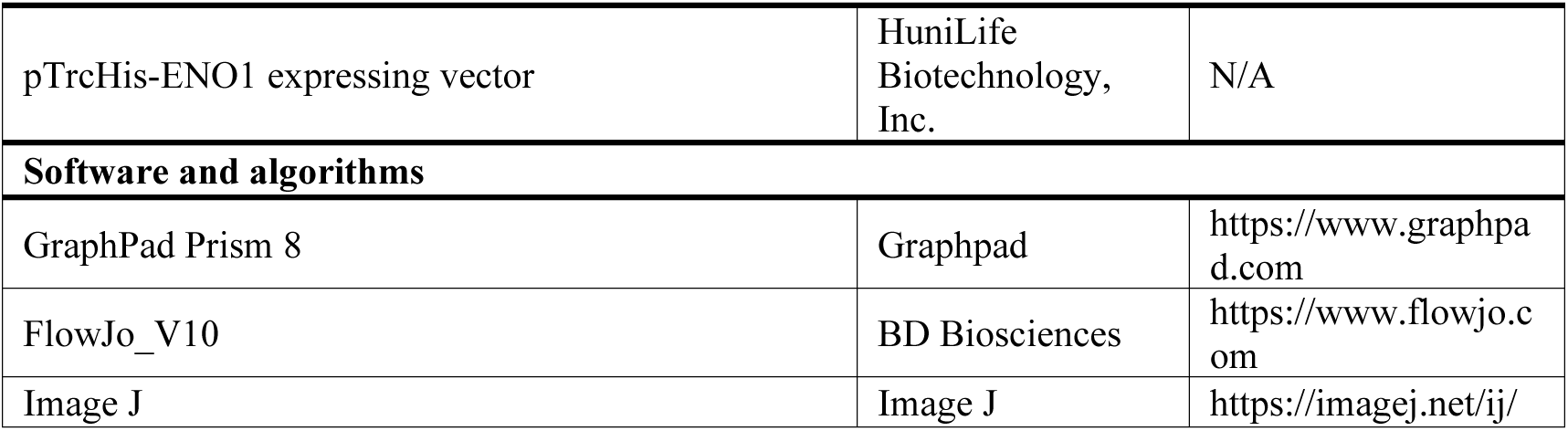
SKEY RESOURCES TABLE.

## EXPERIMENTAL MODEL AND STUDY PARTICIPANT DETAILS

### Cell lines

Murine T lymphoblast EL4 cells and murine alveolar macrophage MH-S cells were cultured in RPMI-1640 medium (Cytiva, MA, USA) supplemented with 10% fetal bovine serum (FBS) (Peak Serum, FL, USA), penicillin-streptomycin-amphotericin B (Biological Industries, Israel), and 0.05 mM β-mercaptoethanol (Cyrusbioscience, Taiwan). All cells were maintained at 37°C in a humidified atmosphere with 5% CO_2_.

### Mice

Male C57BL/6 mice (6-8 weeks old) were purchased from the National Laboratory Animal Center (Taiwan). The mice were housed under standard 12-hour light/dark cycle conditions in rooms with a controlled temperature (22 ± 2°C) and humidity (50 ± 10%). All animal studies were approved by the Institutional Animal Care and Use Committee of National Taiwan University (Approval number: NTU-113-EL-00011).

## METHOD DETAILS

### Isolation of mouse neutrophils from bone marrow and peripheral blood

Mouse bone marrow neutrophils were isolated from the femurs and tibias of C57BL/6 mice using a modified procedure described previously ^45^. Briefly, bone marrow cells were flushed from the bones using Dulbecco’s phosphate-buffered saline (DPBS,

Corning, NY, USA) containing 2% fetal bovine serum (FBS) and 5 mM EDTA. Neutrophils were purified by Percoll (62.5%, Cytiva) density gradient centrifugation at 1,000 × g for 30 minutes. The neutrophil-enriched fraction beneath the 62.5% Percoll layer was collected, washed twice with DPBS, and treated with ammonium-chloride-potassium (ACK) lysis buffer to remove residual erythrocytes. For stimulation experiments, purified bone marrow neutrophils were incubated with lipopolysaccharide (LPS, 100 ng/mL, Sigma-Aldrich, MO, USA) for 3 hours and analyzed for surface ENO1 expression. For ENO1 mRNA and protein expression analysis, mice were challenged intraperitoneally (i.p.) with LPS (2 mg/mL), and bone marrow and peripheral blood neutrophils were isolated by Ly6G⁺ positive selection (MojoSort™ Magnetic Cell Separation, BioLegend, CA, USA) and pooled together.

### Flow Cytometry

For neutrophil and monocyte analysis, single-cell suspensions were first incubated with Fc blocking buffer (containing mAb 2.4G2) for 15 minutes at 4°C, followed by incubation with PE/Cyanine7-anti-CD11b (BioLegend), FITC-anti-Ly6G (BioLegend), and biotin-anti-Ly6B.2 (Bio-Rad, CA, USA) or biotin-anti-Ly6C (BioLegend) for 30 minutes at 4°C. After washing, the cells were incubated with APC-streptavidin (BioLegend) for 30 minutes at 4°C. Dead cells were excluded using 7-AAD staining. For cell surface ENO1 expression analysis, single-cell suspensions were first blocked with mouse serum for 15 minutes at 4°C, followed by incubation with rat-anti mouse ENO1 mAb 7E5 (provided by HuniLife Biotechnology, Taiwan) ^35^ and PE-goat anti-rat IgG (BioLegend) for 30 minutes at 4°C. Afterward, cells were stained with the respective cell surface markers. For NETotic neutrophil analysis, single-cell suspensions were incubated with Fc blocking buffer for 15 minutes at 4°C, followed by incubation with anti-citH3 antibody (Abcam, UK), PE/Cyanine7-anti-CD11b, and FITC-anti-Ly6G for 60 minutes at 4°C. Cells were then incubated with PE-donkey anti-rabbit IgG (BioLegend) for 30 minutes at 4°C. Samples were analyzed on the FACSCanto II flow cytometer (BD Biosciences, NJ, USA), and data, including median fluorescence intensity (MFI), were analyzed using FlowJo software (BD Biosciences).

### RNA extraction and real-time quantitative reverse transcription PCR (RT-qPCR)

Total RNA was purified from cell samples using Qiazol reagent (Qiagen, Germany) following the manufacturer’s instructions. Reverse transcription was performed using the Deoxy+ HiSpec Reverse Transcriptase kit (Yeastern Biotech, Taiwan). RT-qPCR was carried out using the iQ SYBR Green Supermix (Bio-Rad) on a CFX Connect thermal cycler (Bio-Rad) according to the manufacturer’s protocol. mRNA expression levels were normalized to *Actb* (β-actin) and calculated using the 2^−ΔΔCT^ method. The primers used in this study were as follows: For *Eno1*, forward 5’-AGCGATCCTACTGCCAGAAAT-3’ and reverse 5’-GATCGACCTCAACAGTGGGA-3’; for *Actb*, forward 5’-AGGTGTGCACCTTTTATTGGTCTCAA-3’ and reverse 5’-TGTATGAAGGTTTGGTCTCCCT-3’.

### Total cell lysate preparation and Western blotting

Purified neutrophils were resuspended in radioimmunoprecipitation assay (RIPA) buffer (50 mM Tris-HCl, pH 8.0, 150 mM NaCl, 1% Triton X-100, 0.5% SDS) supplemented with a protease inhibitor cocktail (Sigma-Aldrich) and incubated on ice for 30 minutes. After centrifugation at 14,000 × g for 10 minutes, the supernatant containing whole-cell lysates (20 μg of protein) was separated by 10% SDS-PAGE and transferred onto polyvinylidene difluoride (PVDF) membranes (Millipore, MA, USA). The membranes were blocked with Tris-buffered saline containing 0.1% Tween-20 (Cyrusbioscience) (TBST) and 2% BSA (BioVision, CA, USA) for 1 hour at room temperature (RT). They were then incubated overnight at 4°C with primary antibodies against ENO1 (Abcam, UK) or β-actin (Santa Cruz, CA, USA). After extensive washing with TBST, the membranes were incubated with horseradish peroxidase (HRP)-conjugated secondary antibodies (Cytiva) for 1 hour at RT. Protein signals were detected using the T-Pro LumiLong Plus Chemiluminescent Substrate Kit (T-Pro Biotechnology, Taiwan) and visualized using an imaging system. Densitometric analysis of band intensities was performed using ImageJ software (National Institutes of Health, USA).

### Expression of recombinant ENO1 (rENO1)

The cDNA encoding human *ENO1* was cloned into the pTrcHis vector (provided by HuniLife Biotechnology) and expressed in *E. coli* BL21(DE3) cells. Bacterial cultures were incubated at 37°C on an orbital shaker at 140 rpm. When the optical density at 600 nm (A₆₀₀) reached 0.6–0.7, expression of r ENO1 was induced by adding isopropyl-1-thio-β-D-galactopyranoside (IPTG, Sigma-Aldrich) to a final concentration of 0.7 mM, followed by incubation at 25°C for 16 hours. Cells were harvested by centrifugation, resuspended in binding buffer (20 mM NaH₂PO₄, 0.5 M NaCl, 5 mM imidazole, pH 7.0) supplemented with a protease inhibitor cocktail (Sigma-Aldrich), and lysed using a Continuous Flow Cell Disruptor (Constant Systems, UK). rENO1 protein was purified using a HisTrap FF Crude histidine-tagged protein purification column (Cytiva) according to the manufacturer’s instructions. Protein purity was assessed by SDS-PAGE and Western blotting. The purified protein was desalted using a HiTrap Desalting column (Cytiva), and endotoxin was removed using Detoxi-Gel™ Endotoxin Removing Gel (Thermo Fisher Scientific) per the manufacturer’s protocol. The endotoxin level was measured using the ToxinSensor™ Single Test Kit (GenScript, NJ, USA) and determined to be < 0.03 EU/mL.

### Stimulation of MH-S cells

MH-S cells (1 × 10^4^) were seeded in 96-well plates and cultured overnight. The following day, cells were stimulated with rENO1 (50 μg/mL) either alone or in the presence of 7E5 mAb or control IgG. To prevent potential LPS contamination in rENO1, polymyxin B (10 μg/mL) was added. After 20 hours of stimulation, TNF and IL-6 levels in the culture supernatants were measured using ELISA kits (BioLegend).

### Dot blot analysis

The interaction between ENO1 and PLG was assessed using dot blotting. Recombinant ENO1, PLG (Sigma-Aldrich), and BSA (1 μg each) were spotted onto PVDF membrane strips. The membranes were blocked with TBST supplemented with 2% BSA for 1 hour at room temperature. Subsequently, the membranes were incubated with 1 μg/ml of either ENO1 or PLG in PBS for 1 hour at RT. After washing with TBST, immunoblotting (IB) was performed using rabbit anti-ENO1 antibodies (Abcam) or biotinylated goat anti-PLG antibodies (Abcam) for 1 hour at RT. Following additional washes, the membranes were incubated with HRP-conjugated secondary antibodies (Cytiva) or HRP-conjugated avidin (BioLegend), and signals were detected using chemiluminescence as described above.

### ELISA-based protein binding assay

To evaluate the inhibitory effect of 7E5 mAb on the ENO1-PLG interaction, ELISA plates (96-well, Corning) were coated with 10 μg/ml of recombinant ENO1 in PBS and incubated overnight at 4°C. After blocking with ELISA diluent (eBioscience, CA, USA) for 1 hour at RT, the wells were incubated for 2 hours at RT with ELISA diluent containing varying concentrations of 7E5 mAb or a rat IgG control (SouthernBiotech, AL, USA). Subsequently, PLG (10 μg/ml in ELISA diluent) was added and incubated for 2 hours at RT. The wells were then probed with biotinylated anti-PLG antibodies (Abcam) for 1 hour, followed by incubation with HRP-conjugated avidin (BioLegend) for 30 minutes at RT. Colorimetric detection was performed using TMB substrate (BioLegend), and absorbance was measured at 650 nm using a microplate reader (Thermo Fisher Scientific).

### LPS-induced ALI mouse model

To induce ALI and inflammation, mice were i.t. challenged with LPS (2 mg/kg in 50 μL of DPBS) or DPBS alone (sham). To evaluate the effect of anti-ENO1 antibody treatment, mice received an i.v. injection of mAb 7E5 or a rat IgG control (5 mg/kg) 2 hours before LPS challenge. To assess the effect of TXA treatment, mice were administered an i.v. injection of TXA (100 mg/kg) or PBS 0.5 hour before LPS challenge. At 6, 12, or 24 hours post-LPS challenge, blood was collected, mice were sacrificed, BALF was harvested, and lung tissues were prepared for histological analysis.

### Bronchoalveolar lavage and BALF analysis

Bronchoalveolar lavage was performed as previously described ^46^. Briefly, 0.5 mL of PBS containing 5 mM EDTA was instilled into the trachea four times using a 24G i.v. catheter (Terumo, Japan), and BALF was collected. The BALF was centrifuged at 350 × g for 5 minutes, and the supernatant was stored at –20°C for further analysis. The cell pellet was resuspended in PBS, and total cell counts were determined using a hemocytometer. BALF cells were further analyzed by flow cytometry. Cytokine and chemokine concentrations in the BALF were measured using ELISA kits for TNF, IL-6, and keratinocyte chemoattractant (KC) (PeproTech, NJ). Total protein and dsDNA levels in the BALF were quantified using the Pierce™ BCA Protein Assay Kit (Thermo Fisher Scientific) and the Qubit™ dsDNA Quantification Assay Kit (Thermo Fisher Scientific), respectively.

### Measurement of soluble ENO1 by ELISA

Ninety-six-well plates were coated overnight at 4°C with humanized anti-ENO1 mAb HL217 (HuniLife Biotechnology) in PBS. The plates were then blocked with a blocking buffer (Abcam) for 1 hour at RT. Serially diluted recombinant ENO1 (used as the standard) and mouse plasma samples were added and incubated for 2 hours at RT. Next, the wells were incubated with 7E5 mAb (1 μg/ml in blocking buffer) for 1 hour at RT, followed by incubation with biotinylated mouse anti-rat IgG (Thermo Fisher Scientific) for 1 hour at RT. Avidin-HRP (BioLegend) was then added and incubated for 30 minutes at RT. The reaction was visualized using TMB substrate solution (BioLegend), and absorbance was measured at 650 nm using a microplate reader.

### Neutrophil transmigration assay

Neutrophil transmigration was assessed using a Transwell system, as previously described ^47^. Briefly, Transwell inserts (6.5-mm diameter, 3.0-μm pores; Corning) were coated with ECM gel (200 μg/mL; Sigma-Aldrich) and incubated overnight at 37°C.

Purified neutrophils from naïve mice were resuspended in RPMI-1640 medium at 3 × 10^6^ cells/mL and preincubated with 10 μg/mL 7E5 mAb, rat IgG control, or 10 mM TXA for 30 minutes at 37°C. The cells were then incubated with 500 nM Glu-plasminogen (Glu-PLG; Technoclone, Austria) for 1 hour at 37°C. After washing to remove unbound PLG, neutrophils were resuspended in RPMI-1640 medium containing 50 U/mL urokinase (Taiwan Green Cross, Taiwan), and 3 × 10^5^ cells were loaded into the Transwell inserts. The lower chambers were filled with 600 μL RPMI-1640 medium with or without 50 ng/mL CXCL1. After incubation at 37°C for 18 hours, migrated cells in the lower chambers were counted using a hemocytometer.

### Necrotic cell-induced peritonitis mouse model

Necrotic EL4 cells were prepared using the heat-shock method as previously described ^48^. To induce sterile peritonitis, C57BL/6 mice were injected i.p. with 2 × 10^7^ necrotic EL4 cells in 0.5 mL PBS, while control mice received 0.5 mL PBS alone. To assess the effect of anti-ENO1 antibody treatment, mice were administered i.p. with 5 mg/kg of 7E5 mAb or rat control IgG 2 hours before necrotic cell challenge. Twelve hours post-challenge, blood was collected, mice were euthanized, and peritoneal lavage was performed using PBS supplemented with 2% FBS and 5 mM EDTA. PECs were counted using a hemocytometer and analyzed by flow cytometry.

### Histopathology

Mouse lung tissues were perfused and fixed in 10% formalin, then embedded in paraffin and stained with hematoxylin and eosin (H&E) at the Histology Center, Institute of Molecular and Comparative Pathobiology, National Taiwan University. Lung injury was assessed by a veterinarian using a previously described scoring system ^34^. Briefly, lung sections were graded on a scale of 0–2 based on the following parameters: (A) neutrophils in the alveolar space, (B) neutrophils in the interstitial space, (C) proteinaceous debris filling the airspaces, and (D) alveolar septal thickening. For each sample, twenty high-magnification fields were randomly selected, and the total lung injury score was calculated using the formula: [(20 × A) + (14 × B) + (7 × C) + (2 × D)] / 20 × 72).

### Statistics

Statistical analyses were performed using GraphPad Prism (version 8.0). Data are presented as the mean ± standard error of the mean (SEM). Group comparisons were conducted using an unpaired Student’s *t*-test for two-group comparisons and a one-way ANOVA followed by Tukey’s post hoc test for multiple-group comparisons. Statistical significance was set at *p* < 0.05.

